# The role of visual and olfactory cues in social decisions of guppies and zebrafish

**DOI:** 10.1101/2021.05.29.446265

**Authors:** Maria Santacà, Marco Dadda, Angelo Bisazza

**Author notes:** Corresponding author: Maria Santacà. Department of General Psychology, Via Venezia 8, 35131 Padova (Italy)., Co-authors correspondence: Marco Dadda; Angelo Bisazza.

## Abstract

Vision and olfaction are expensive to maintain, and in many taxa there appears to be a trade-off in investment between the two sensory systems. Previous work has suggested that guppies (*Poecilia reticulata*) and zebrafish (*Danio rerio*) may differ in the relative importance they place on these two senses in social interactions. In this study, we directly examined this issue by experimentally contrasting olfactory and visual information in social situations. In the first experiment, we found that guppies spent more time where conspecifics were visible than where they could smell them. On the contrary, zebrafish spent significantly more time in an empty compartment containing smell of conspecifics than in a compartment in which they only saw them. The difference was not large, suggesting that both species integrate various types of information to locate a nearby shoal. In two subsequent experiments, we studied the role of vision and smell in the discrimination of the quality of the social group, namely the number and the familiarity of its members. Zebrafish and guppies were confirmed to rely on different senses to estimate the size of a social group, whereas they did not differ in the discrimination of familiar and non-familiar conspecifics which appears to be based equally on the two senses. Similarly to what happens in other vertebrate clades, we suggest that, among teleosts, there are large differences in the relative importance of the different senses in the perception of the external world.

## INTRODUCTION

Three main senses—vision, olfaction, and hearing—allow animals to collect information about distant objects or events. Two of these senses are particularly costly for the organism as indicated by a considerable metabolic maintenance cost (up to 15% of resting metabolism in some species) for the visual system and by a large proportion of genes (up to 5% of the genome) coding for proteins involved in the olfactory system (e.g., Godfrey, Malnic, & Buck, 2004; Moran, Softley, & Warrant, 2015; Wong-Riley, 2010). This frequently determines an evolutionary trade-off between vision and olfaction, and though vertebrates share essentially the same sensory systems, there is considerable variation in the relative importance of these senses in the different taxa (e.g., Barton, Purvis, & Harvey, 1995; Kotrschal, Van Staaden, & Huber, 1998).

Some mammalian orders (e.g., carnivorans and rodents) are composed of “macrosmatic” species, which primarily rely on hearing and chemical information for important functions such as foraging, antipredator responses, and social and sex interactions (Moulton, 1967). Conversely, haplorhine primates (monkeys and apes) are microsmatic and they preferentially use their sophisticated visual systems for the same functions (Niimura, Matsui, & Touhara, 2018; Smith, Rossie, & Bhatnagar, 2007). Birds appear more homogeneous in this respect, and with the exception of a handful of species, they are all microsmatic (Stager, 1967). There is no such systematic information for the other vertebrate classes. Reptiles are usually classified as macrosmatic although some lizards are microsmatic and possess an advanced visual system (Goldby & Gamble, 1957). Amphibians are well known to use vision for prey capture. The importance of chemical senses is less investigated but there is evidence that they use the sense of smell for some functions such as mating and habitat selection (e.g., Ewert, 1987; Houck, 2009; Martof, 1962).

Due to the physics of sound propagation in water, for small aquatic species, hearing is effective for social communication but much less effective for determining the direction from which a signal comes (Nummela & Thewissen, 2008). In teleosts, this leaves smell and vision a prominent role in locating and recognizing objects that are far from the animal’s body. As it occurs in air, in water too these senses are subject to various constraints. For example, chemical senses are constrained by the direction of water flow. Conversely, vision can be impeded by the presence of obstacles or by the turbidity of water (e.g., Finelli, Pentcheff, Zimmer-Faust, & Wethey, 1999; Lunt & Smee, 2015; Sundin, Berglund, & Rosenqvist, 2010).

There is scarce information about the relative importance of the different sensory modalities in fish. Nocturnal species unsurprisingly rely more on tactile, chemical and electric senses than on vision for feeding, mating and orientation (e.g., Atta, 2013; Teichmann, 1954). Species that live in bright light conditions are generally considered microsmatic; however, data from the anatomy and physiology of sensory systems suggest that even among diurnal species, there might be a wide variability in the relative importance of smell and vision (e.g., Hara, 1975; Kasumyan, 2004; Teichmann, 1954).

A recent study examined the efficiency of four teleost species in detouring a transparent barrier to reach a goal (a group of conspecifics) placed behind it (Santacà, Busatta, Lucon-Xiccato, & Bisazza, 2019). While the performance of three species (*Poecilia reticulata, Xenotoca eiseni, Oryzias sarasinorum*) was comparable to that shown on average by mammals and birds, one species, the zebrafish (*Danio rerio*), showed a proficiency level in this test that was previously observed only in corvids and apes. Control experiments suggested that zebrafish enjoyed a sensory advantage over the other species, in that it was guided by olfactory cues (to which the barrier was opaque) to head to the social stimuli. The hypothesis that zebrafish have a more sophisticated olfactory system and may thus rely primarily on olfaction for sensing conspecifics, is supported by the comparison of the microstructure of central and peripheral olfactory systems of guppy and zebrafish (Bettini, Lazzari, Ciani, & Franceschini, 2009; Lazzari, Bettini, & Franceschini, 2014).

Fish have complex cognitive functions that they also use in social contexts. These allow, for example, recognition of a fish that has previously cooperated with them when inspecting a predator, to copy the mate choices of other individuals or learn new traditions from experienced conspecifics (reviewed in Bisazza, 2010; Bshary, Wickler, & Fricke, 2002). Two aspects of social cognition have been thoroughly investigated, the ability to estimate the numerosity of a social group and the ability to discriminate between familiar and unfamiliar individuals. All the gregarious fishes studied so far show a strong tendency to join the group that contains the greatest number of conspecifics, a behavior believed to be associated with anti-predatory advantages (e.g., Agrillo & Bisazza, 2018; Agrillo, Miletto Petrazzini, & Bisazza, 2017; Krause, Ruxton, Ruxton, & Ruxton, 2002). In most studies, fish were allowed to use only visual information, and in a few others visual and olfactory information were used together. However, no study investigated the relative importance of the two senses in this context. Teleosts are also able to discriminate conspecifics with which they have lived from unfamiliar ones. Some studies have highlighted the role of vision, demonstrating, for example, the ability to recognize the face of family members; others have investigated the role of the chemical senses (e.g., Courtenay, Quinn, Dupuis, Groot, & Larkin, 2001; Wang & Takeuchi, 2017; Ward & Hart, 2003). The only study investigating the relative importance of these two senses, found that fathead minnows (*Pimephales promelas*) recognized familiars in the presence of olfactory cues alone or olfactory plus visual cues, but not when visual cues alone were available (Brown & Smith, 1994).

In this study, we carried out three experiments to verify whether guppies and zebrafish differ in the relative importance of vision and smell in their social decisions. In the first experiment, individuals searching for conspecifics were allowed to choose between a compartment in which they could only smell a group of conspecifics and one in which the social group was only visible. The second experiment was on shoal size discrimination. On one side, the subjects saw six conspecifics and perceived the smell of three, while on the other side, they saw three conspecifics but perceived the smell of a group of six. In the third experiment, we studied the role of chemical and visual communication in the recognition of familiar fish, also by dissociating the visual and chemical information.

## METHODS

### Experiment 1: The role of vision and olfaction in social group location

#### Subjects

Twelve zebrafish and 12 guppies were tested in this experiment. All fish were experimentally naïve. Twenty-eight additional fish for each species were used as visual stimuli or as donors of chemical cues. Only adult females were used as subjects or stimuli, to avoid the confound of sex. Zebrafish were bought from a local pet shop two months before the experiment whereas the guppies originated from some stocks collected from a large (>10,000 fish) self-sustaining population of an artificial pond. These fish descend from wild ancestors that were collected in the lower Tacarigua River (Trinidad) in 2002 and were brought in the laboratory two months before the experiment. In our laboratory, the two species were kept separated and maintained in 400-L opaque plastic tanks enriched with natural plants and gravel bottoms. The tanks were also provided with two biomechanical filters and 30-w fluorescent lamps (12h:12h light/dark photoperiod). The water temperature was constant at 26 ± 1 °C. Both species were fed twice a day: they were fed with commercial food flakes (Aqua tropical, Padovan®) in the morning and live *Artemia salina* nauplii in the afternoon.

#### Apparatus

The test apparatus (Figure 1) consisted of five tanks of different sizes, a central subject tank, two tanks for visual stimuli and two for olfactory stimuli. The subject tank (60 × 40 × 32 cm) was internally shaped as an hourglass by means of a transparent plastic structure. On the bottom of the tank, a hole allowed the rapid and complete emptying after each trial. This water was filtered by activated carbon and biomechanical filters and stocked in a 400-L reservoir to be reused. The subject tank was not directly lit but received indirect light from the two visual stimuli tanks. A video recorder was placed above the subject tank. To become familiar with the shape of test apparatus, before the experiments the 12 subjects were kept together in two identical habituation tanks, one for each species. The habituation tank (60 × 40 × 32 cm) replicated the test tank in shape and size but was enriched with vegetation, water filtering mechanisms and a gravel bottom, similar to the maintenance tanks. Each tank was also provided with a heater, to maintain a constant water temperature at 25 ± 1 °C, and two 15 W fluorescent lamps (12h:12h light/dark photoperiod).

**Figure 1.**
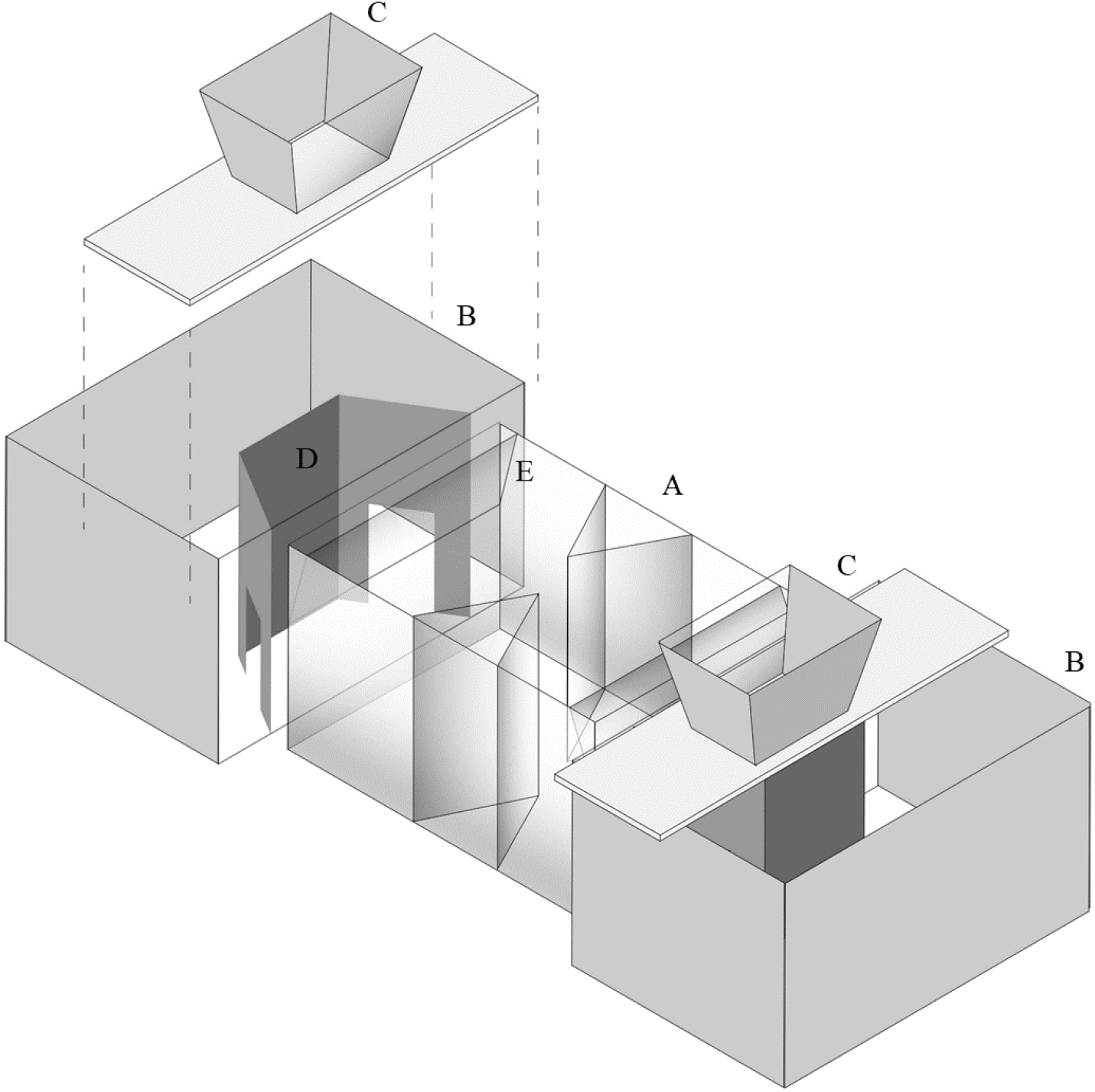
Experimental apparatus. The experimental apparatus was composed of the subject tank (A), two visual stimuli tanks (B) and two olfactory stimuli tanks (C). The visual stimuli tanks had a visible compartment (D) that housed the visual stimuli during the experiments. The olfactory stimuli tanks were placed above the visual stimuli tanks. Olfactory cues were released along the short wall in a transparent structure (E) that served to slow the diffusion of odour.

**Figure 2.**
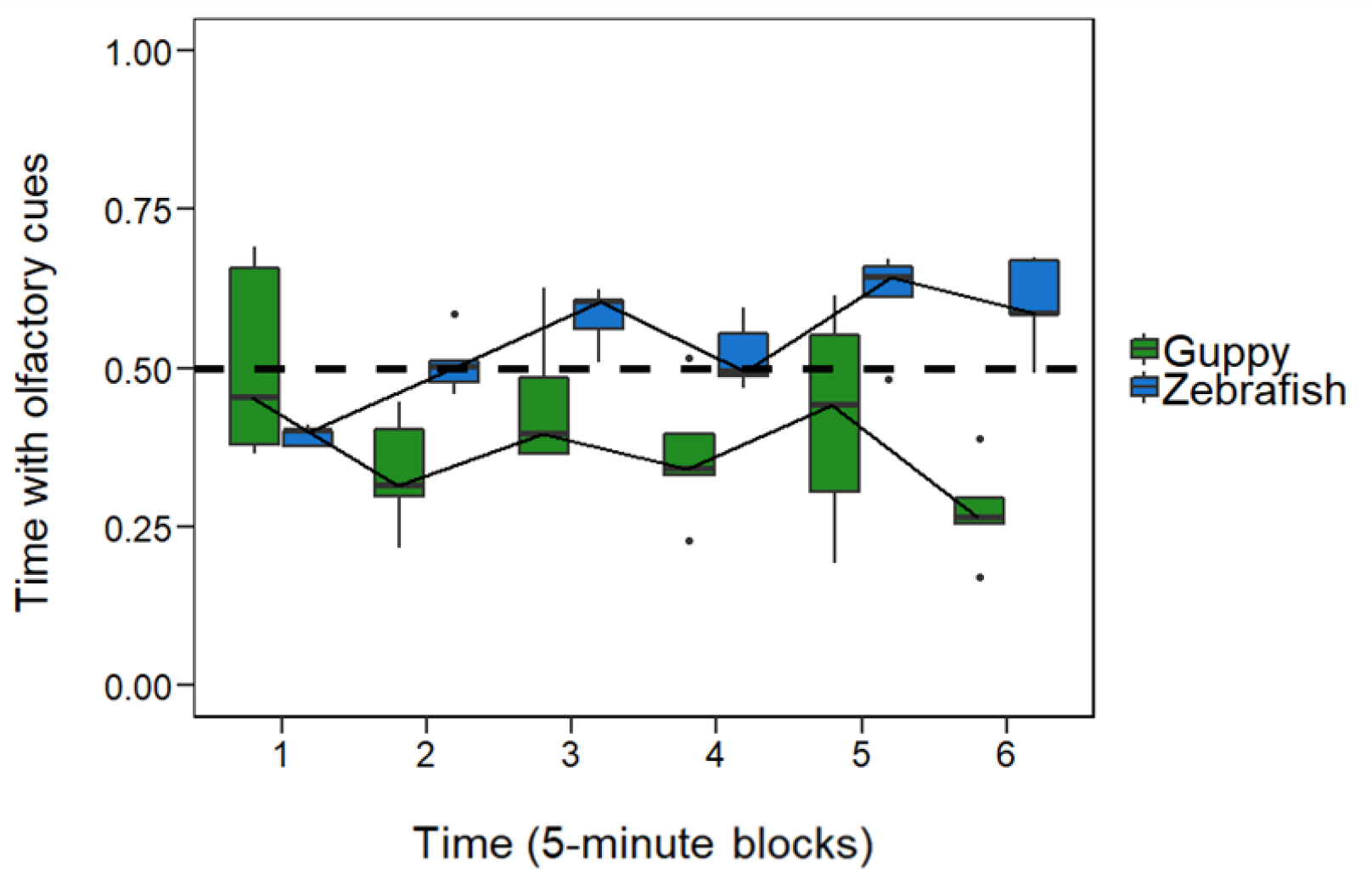
Results of the first experiment. Comparison of the mean proportion of time spent in the compartment with olfactory cues between guppies and zebrafish. The boxplots report median, first quartile, third quartile, ranges, and outliers (data points 1.5 interquartile ranges smaller than the first quartile or greater than the third quartile).

**Figure 3.**
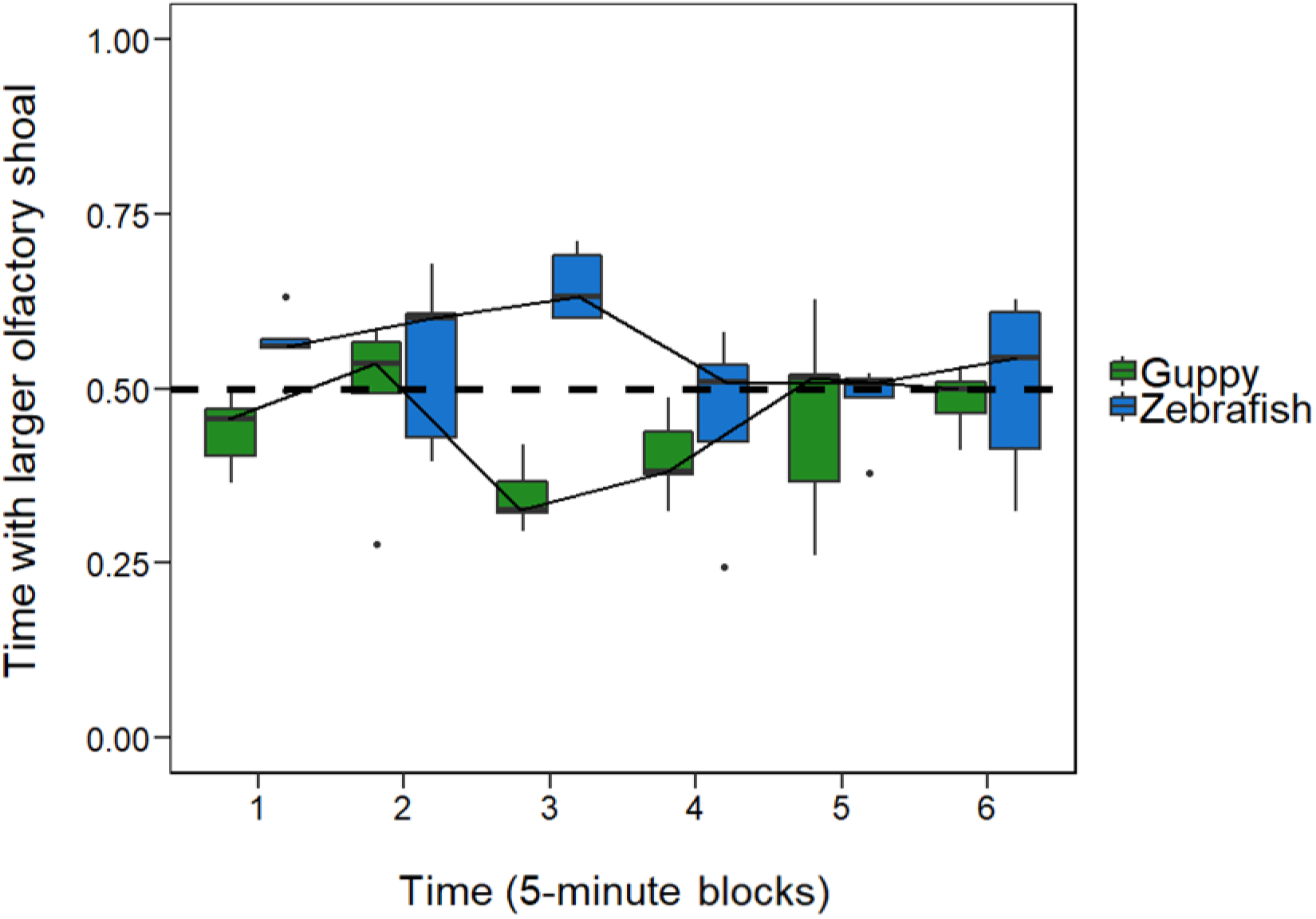
Results of the second experiment. Comparison of the mean proportion of time spent in the compartment with of the larger olfactory shoal between guppies and zebrafish. The boxplots report median, first quartile, third quartile, ranges, and outliers (data points 1.5 interquartile ranges smaller than the first quartile or greater than the third quartile).

**Figure 4.**
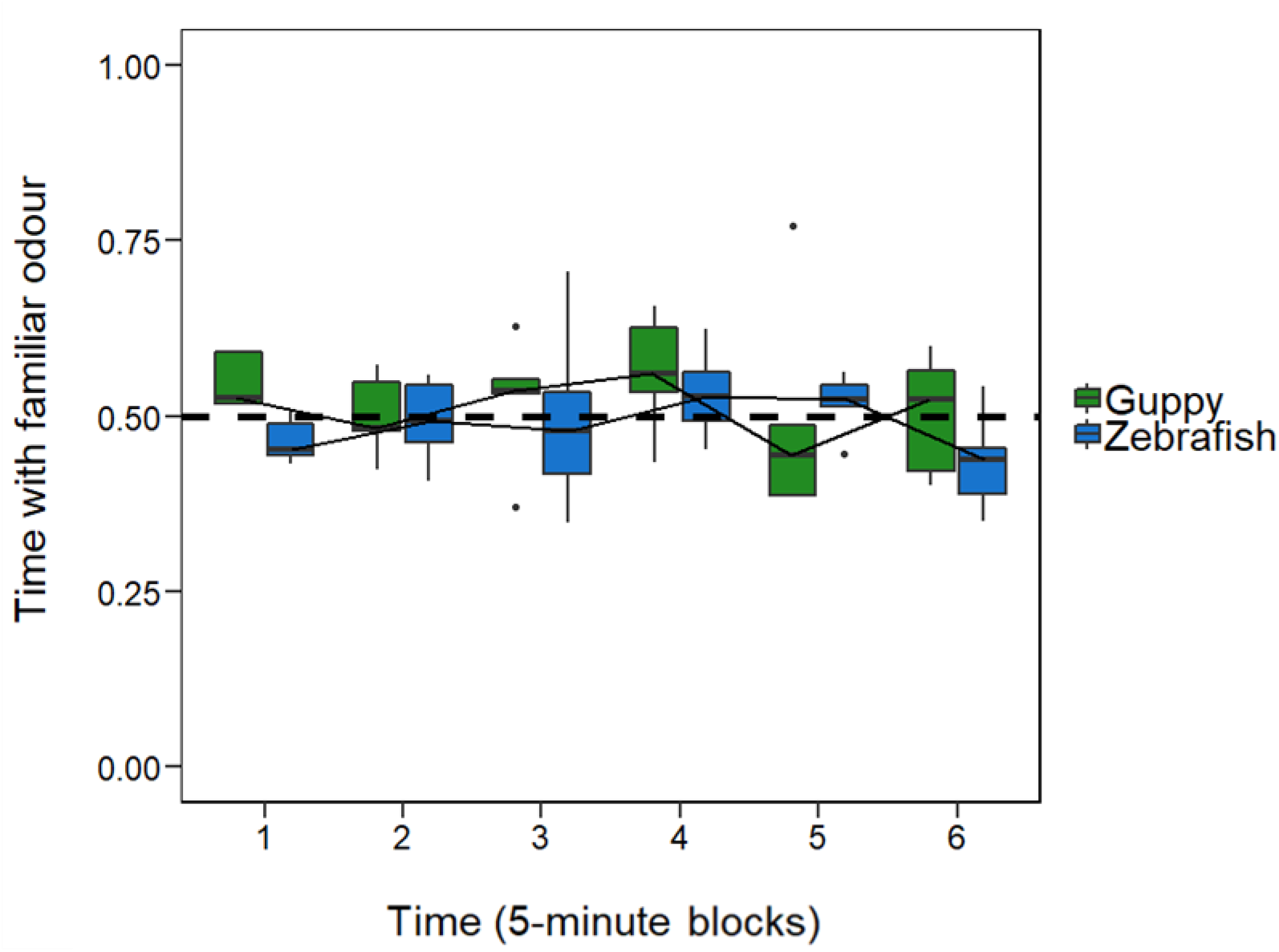
Results of the third experiment. Comparison of the mean proportion of time spent in the compartment with familiar odour between guppies and zebrafish. The boxplots report median, first quartile, third quartile, ranges, and outliers (data points 1.5 interquartile ranges smaller than the first quartile or greater than the third quartile).

Each visual stimuli tank (60 × 40 × 32 cm) was divided in two compartments: a smaller front compartment visible to the subject and a larger back compartment containing a filter, natural vegetation and a gravel bottom. The two compartments were separated by means of two sliding transparent panels (8 × 5 cm). To improve the visibility of stimuli, a white panel served as a background of the visible compartment. During the seven days of the stimuli habituation, the fish were free to move between the two compartments, but they were fed only in the visible compartment. Eight visual stimuli were inserted one week before the beginning of the experiment and lived in the stimuli tanks for the duration of the experiment. They were free to move between the two compartments but were fed only in the visible compartment.

Each olfactory stimuli tank (28 × 19 × 14 cm) was placed above a visual stimuli tank and lit by a 15-w fluorescent lamp. It was externally covered with green plastic and not visible from the subject tank. A transparent small pipe, controlled by a valve, released the water in the middle of the short wall of the subject tank. When the valves were turned on, each tube delivered water at a rate of 3 ml/min. A transparent acrylic sheet (40 × 18 cm) placed along the short wall served to slow the diffusion of odour in the subject compartment. In preliminary tests using dye, we observed that water diffusion was slow and that the dye did not reach the corridor within the first 30 minutes (the duration of the test).

#### Procedure

A week before the start of the experiment, subjects were moved to an 80-L habituation tank. The four stimulus tanks were set up 90 min before the start of the experiment. On one side, six visual stimuli were confined in the front compartment of their tank, whereas on the other side, no visual stimulus was visible. Four conspecifics were inserted in one olfactory stimuli tank, whereas the other contained no fish. Fish used as stimuli came from a separate tank and were not familiar to the subjects.

The subject tank was filled with 30 cm of water from the 400-L reservoir by means of a pump. Five minutes before the start of a trial, the valve of the two olfactory tanks was open, and the odour released in the subject tank. At the beginning of the trial, a subject was moved from the habituation tank, released in the middle of the subject tank and observed for 30 minutes. In one side of the tank, the subject could see the conspecifics with no smell. In the other side, it could smell conspecific odour but did not see conspecifics. The side in which fish could see or smell the conspecifics was the same during an experimental day, but it was reversed each day. At the end of the trial, the subject was removed, released in a post-experimental tank and used for reproduction and the tank was completed emptied. The subject tank and the olfactory tanks were then filled with new aged water.

Based on the video recordings, we scored the performance of the subjects. We recorded whether the subject spent more time in the compartment with the visual stimuli or in the compartment with the olfactory cues. We used the computer software originally developed in our laboratory (‘Ciclic Timer’, written in Delphi 5 Borland). We also counted the number of passages between the two compartments. Two different experimenters analysed one third of the videos of both species to assess inter-rater reliability.

#### Statistical Analysis

Analyses were performed in R version 3.5.3 (The R Foundation for Statistical Computing, Vienna, Austria, http://www.r-project.org). We assessed inter-rater reliability between two experimenters: we tested for a correlation between the two scorers using the Spearman’s rank method. As inter-rater reliability was excellent (*ρ* = 0.979, *P* < 0.001), we conducted the analyses using the database of only one experimenter. We analysed the subjects’ performances (the time spent in each area after log transformation due to right-skewed distribution) with linear mixed-effects models (LMMs, ‘lmer’ function of the ‘lme4’ R package). To assess whether the subjects spent more time in a compartment compared to the other and whether the two species differed, we fitted the model with the compartment and the species as fixed effects. In addition, we fitted the model with 5-minutes blocks as a fixed effect to assess whether the subjects’ performances changed across time. In addition, the individual ID was fitted as a random effect. Subsequently, all pairwise comparisons were performed with Tukey post-hoc tests.

#### Ethical Note

Experiments were conducted in compliance with the law of the country (Italy) in which they were performed (Decreto legislativo 4 marzo 2014, n. 26). The experimental procedures were approved by the Ethical Committee of Università di Padova (protocol n. 16/2018). Subjects did not appear to be stressed during the experiments; after the test, they were released in a tank and used only for breeding.

### Experiment 2: The role of vision and olfaction in shoal size assessment

#### Subjects, apparatus and procedure

In Experiment 2, 12 adult female zebrafish and 12 adult female guppies were tested. Zebrafish and guppies were tested in the same apparatus and with the same procedure as the previous experiment. However, in this case, they could see six conspecifics and smell the odour of three conspecifics in one compartment, whereas in the other one, they could see three conspecifics and smell the odour of six conspecifics.

#### Statistical Analysis

Analyses were performed in R version 3.5.3 (The R Foundation for Statistical Computing, Vienna, Austria, http://www.r-project.org). We assessed inter-rater reliability as Experiment 1: the inter-rater reliability was also excellent in this experiment (*ρ* = 0.982, *P* < 0.001). We analysed the subjects’ performance with LMM as in the previous experiment, fitting the model with the compartment, the species and 5-minute blocks as fixed effects. Individual ID was fitted as random effect. All pairwise comparisons were performed with Tukey post-hoc tests.

### Experiment 3: The role of vision and olfaction in familiar conspecifics recognition

#### Subjects, apparatus and procedure

In Experiment 3, 12 adult female zebrafish and 12 adult female guppies were tested. Zebrafish and guppies were tested in the same apparatus and with the same procedure as the previous experiments. In one compartment they could see four familiar conspecifics and smell the odour of four unfamiliar conspecifics whereas in the other compartment they could see four unfamiliar conspecifics and smell the odour of four familiar conspecifics. In this experiment, the pre-experimental treatment was modified. To acquire familiarity, experimental subjects and stimulus fish were maintained together for 14 days (Barber & Wright, 2001; Griffiths & Magurran, 1997). Prior to the experiment, the fish were subdivided in six familiarization tanks (50 × 20 × 35 cm), eight fish in each tank. Internally, each tank was identical to the visible compartment of the visual stimuli tanks. At the end of this period, four fish were randomly selected to become stimuli; two were selected to become subjects; and two other fish remained in the familiarization tank as companion fish to avoid subjects remaining isolated prior to the test. During each experimental day, we tested the subjects of two familiarization tanks. Two hours before the experiment, the two groups of four stimulus fish were moved, each to one of the two visual stimuli tanks. The same fish also served as donors for the olfactory stimuli. By means of a system of pumps, the water from each visual stimuli tank circulated in the olfactory stimuli tank on the opposite side of the apparatus, so that, during the experiment, the odour of familiar fish could be released near unfamiliar ones and vice versa.

#### Statistical Analysis

Analyses were performed in R version 3.5.3 (The R Foundation for Statistical Computing, Vienna, Austria, http://www.r-project.org). We assessed inter-rater reliability as all previous experiments. The inter-rater reliability was also excellent in this case (*ρ* = 0.973, *P* < 0.001). As the previous experiments, we analysed the subjects’ performances with LMM. We fitted the model with the compartment, the species and 5-minute blocks as fixed effects, and the individual ID was fitted as a random effect. We have also considered the familiarization tank as fixed effect. All pairwise comparisons were performed with Tukey post-hoc tests.

## RESULTS

### Experiment 1: The role of vision and olfaction in social group location

The average number of passages from one compartment to the other was 54.16 ± 18.09 for zebrafish and 17.00 ± 7.82 for guppies. Zebrafish spent 822.92 ± 108.01 s (mean ± SD; 45.72 ± 6.00 % of the time) in the compartment with the visual cues and 977.00 ± 108.00 s (54.28 ± 6.00 %) in the compartment with the olfactory cues. Guppies spent 1093.00 ± 202.80 s (60.76 ± 11.25 %) in the compartment with the visual cues and 705.67 ± 202.14 s in the compartment with the olfactory cues (39.24 ± 11.25 %).

The LMM on the time spent in the two compartments revealed that there was a significant effect of the compartment (*F*_1,2834_ = 5.671, *P* < 0.05) and of the species (*F*_1,22_ = 29.407, *P* < 0.001) and no significant effect of the time (i.e., considering blocks of 5 minutes; *F*_5,2834_ = 1.290, *P* = 0.265). The interaction compartment × time was significant (*F*_5,2834_ = 2.559, *P* < 0.05) such as the interaction compartment × species (*F*_1,2834_ = 50.115, *P* < 0.001). All the other interactions were not significant (all *P*-values > 0.265). The Tukey post hoc tests revealed that for both species the time spent in a compartment was significantly different from the time spent in the other compartment (all *P*-values < 0.005). In particular, guppies spent significantly more time in the compartment with the visual cues, whereas zebrafish spent significantly more time in the compartment with the olfactory cues suggesting that the two species were using different sensorial information to locate the conspecifics.

### Experiment 2: The role of vision and olfaction in shoal size assessment

The mean number of passages from one compartment to the other was 45.33 ± 11.11 for zebrafish and 15.83 ± 7.71 for guppies. Zebrafish spent 819.92 ± 88.69 s (45.55 ± 4.93 %) in the compartment with the six conspecific visual stimuli and 980.08 ± 88.69 s (54.45 ± 4.93 %) in the compartment with the olfactory cues of six conspecifics. On the contrary, guppies spent 964.75 ± 200.16 s (53.70 ± 11.05 %) in the compartment with the larger visual shoal and 831.42 ± 197.73 s (46.30 ± 11.05 %) in the compartment with the larger olfactory shoal.

The LMM on the time spent in the two compartments revealed that there was a significant effect of the compartment (*F*_1,2834_ = 22.332, *P* < 0.001), of the species (*F*_1,22_ = 11.315, *P* < 0.01) and no significant effect of the time (*F*_5,2834_ = 1.938, *P* = 0.085). No other interactions were significant (all *P*-values > 0.265). The Tukey post hoc tests revealed that, for both species, the time spent in a compartment was significantly different from the time spent in the other compartment (all *P*-values < 0.05). In particular, guppies spent significantly more time in the compartment with the visual cues of the larger shoal, whereas zebrafish significantly spent more time in the compartment with the odour cues of the larger shoal.

### Experiment 3: The role of vision and olfaction in familiar conspecifics recognition

The mean number of passages from one compartment to the other was 56.42 ± 16.49 for zebrafish and 31.83 ± 22.91 for guppies. Zebrafish spent 911.00 ± 135.6 s (50.62 ± 7.52 %) in the compartment with the familiar conspecific visual stimuli and 885.50 ± 135.22 s (49.37 ± 7.52 %) in the compartment with the olfactory cues of familiar conspecifics. Guppies spent 856.75 ± 13.77 s (47.59 ± 7.32 %) in the compartment with the familiar visual stimuli and 943.42 ± 131.74 s (52.41 ± 7.32 %) in the compartment with the familiar olfactory cues.

The LMM on the time spent in the two compartments revealed that there was no significant effect of the compartment (*F*_1,2834_ = 1.287, *P* = 0.257), of the species (*F*_1,12_ = 2.025, *P* = 0.180), of the time (*F*_5,2834_ = 2.166, *P* = 0.055) and of the familiarization tank (*F*_10,12_ = 0.343, *P* = 0.950). The only significant interaction was the interaction compartment × time (*F*_5,2834_ = 3.295, *P* < 0.01). No other interactions were significant (all *P*-values > 0.066).

## DISCUSSION

In the first experiment, we tested whether guppies and zebrafish were more attracted by the sight or by the smell of group of conspecifics when these sensory cues were experimentally dissociated. Based on previous behavioural and anatomical evidence we predicted that zebrafish would rely on olfactory cues more than guppies to locate a group that is close by. Indeed, zebrafish spent more time in the compartment with odour while guppies did the reverse. The difference between the two species is not large (61% with visual cues in guppy and 54% with odour in zebrafish), meaning that both species probably use multiple cues to locate a nearby shoal. Nonetheless, it is remarkable that zebrafish spent significantly more time in a compartment where no fish was visible but where they could smell the presence of conspecifics. In the two subsequent experiments, we investigated whether this sensory difference is limited to the function of locating a nearby shoal or if it extends to other aspects of social interactions such as choosing larger shoals or familiar conspecifics.

When alone and potentially at risk, small social teleosts prefer to join the larger of two shoals, generally showing an extreme accuracy in discriminating the quantity of fish in a group (Agrillo & Bisazza, 2018; Agrillo et al., 2017). There is scarce information about the proximate mechanisms of such discrimination. In the second experiment, which contrasted visual and olfactory information during a shoal size preference test, we found the same sensory difference between the two species that had emerged in the first experiment. Guppies spent more time in the compartment in which they could see six fish but smell three, while zebrafish stayed longer where they could smell six fish and see three. Again, the difference between species was not large (in both species approximately 54% in the preferred compartment), indicating that, while choosing the larger shoal, both species likely integrate visual and olfactory information to increase the accuracy of discrimination. Indeed, the integration of various sources of information to obtain a more accurate number estimation is a typical feature of humans and other vertebrates and it was is commonly observed in fish studies (Agrillo, Piffer, & Bisazza, 2011; Suanda, Tompson, & Brannon, 2008; Tomonaga, 2008).

All teleosts studied so far have been found to prefer familiar to unfamiliar conspecifics. Research on proximate mechanisms investigated either chemical or visual communication in recognizing family members, although the relative importance of the two senses was rarely examined (reviewed in Ward & Hart, 2003). In the third experiment, we examined this issue by dissociating olfactory and visual cues during a shoal choice preference test. Guppies and zebrafish spent a similar amount of time in the compartment with visible familiars or in the compartment with their odour, and we found no difference between the two species. The simplest explanation for our results is that, in neither species, one cue is singly more important than the other for the recognition of familiar fish. Discriminating those conspecifics with which one fish has previously interacted from all other fish of its population is likely a challenging task, dependent upon recognizing subtle individual differences. We can reasonably expect most species to use all available information to increase accuracy in discrimination.

There are, however, other explanations for these results. The first is that, in this particular context, choosing familiar shoal mates is not particularly relevant. The shoal choice test simulates the situation in which a fish is isolated in an unfamiliar, potentially dangerous environment and must therefore rapidly rejoin a group. The advantages usually hypothesized for associating with familiar individuals include reduced aggression, increased reciprocity and more efficient social learning (reviewed in Ward & Hart, 2003). As regards potential anti-predator benefits, groups composed of familiar individuals have been found to show more cohesion, but there is little evidence that fish increase their preferences for familiars under predation hazard (Brown, 2002; Chivers, Brown, & Smith, 1995; Griffiths, 1997). If associating with familiar fish does not confer a large anti-predatory benefit, it could be advantageous to reach the first available group in a risky situation. Hence, selectivity towards familiars is expected to be extremely reduced in our experimental condition.

Another possibility is that there are individual differences and that, when information conflicts, some individuals rely more on smell and others more on vision to recognize familiar fish, with the consequence that the sample, on average, does not evidence a preferred cue. Intraspecific differences in the importance of various senses exist in several vertebrate species (Knott, Berg, Ribot, Endler, & Bennett, 2017; McGreevy Grassi, & Harman, 2004; Wrzesniewski, McCauley, & Rozin, 1999) and have been documented in at least in one fish species, the stickleback, where the importance of smell on social discrimination varies among populations (Hiermes, Mehlis, Rick, & Bakker, 2015). Interestingly, in this experiment, we found a significant compartment × time interaction, which suggests that different cues are preferred at different times of the test. It is possible that individuals differ in the timing they exhibit the different preferences, a factor that could also have contributed to obscuring existing differences in the preferred sensory cue.

What are the explanations for the consistent interspecific differences in the use of different sensory modalities found in this study? As this work examines only two species, there are various possibilities to explain our results. The first hypothesis is that the differences are in relation to specific ecological characteristics of the habitat in which these two species evolved. The transparency of water does not seem a key factor differentiating the habitats. Both species inhabit slow running, shallow, clear steams and are less frequently found in deep and turbid lowland waters. Zebrafish are generally reported to inhabit steams with dense vegetation whereas this seems infrequent in most habitats inhabited by guppies. Although an accurate comparison of the visibility of a conspecific in the habitats of the two species is difficult to make (Engeszer, Patterson, Rao, & Parichy, 2007; Magurran, 2005; Spence et al., 2006). In addition, sensory systems may have diverged for selective factors other than those related to sociality, for instance in relation to the specific predators or preys present in their natural habitats (e.g., Fischer, Oberhummer, Cunha-Saraiva, Gerber, & Taborsky, 2017; Kelley & Magurran, 2003; Tang, Zhou, Chen, & Zheng, 2013).

The second possibility is that the two species belong to two clades that have taken two different evolutionary paths, as happened, for example, for the different orders of mammals. In the study of Santacà et al. (2019), *Xenotoca eiseni* (same order, namely Cyprinodontiformes, of guppy) and *Oryzias sarasinorum* (belonging to the closely related order Beloniformes) tended to be similar to guppies in detouring a transparent barrier. The authors suggested that there may be differences in sensory ecology between the two largest clades of teleosts, Ostariophysi (to which zebrafish belong) and Acanthopterygii (to which guppies belong) which diverged about 220 million years ago. There is a lack of behavioral data to sustain this hypothesis, which seems to be indirectly supported from the comparison of the olfactory system in the different species of teleosts. Based on morphological, physiological or molecular data, some fish have been classified as microsmatic, others as microsmatic. Most species classified as microsmatics on such basis, such as guppy, medaka (*Oryzias latipes*), northern pike (*Esox lucius*), three-spined stickleback (*Gasterosteus aculeatus*), Convict cichlid (*Amatitlania nigrofasciata*), Japanese flyingfish (*Cheilopogon agoo*) and broadnosed pipefish (*Syngnathus typhle*) belong to Acanthopterygii, although sporadic species in this superorder appear to be macrosmatic. In contrast, considered macrosmatic are all examined species belonging to the three large orders of Ostariophysi: the Cypriniformes such as zebrafish, common minnow (*Phoxinus phoxinus*), goldfish (*Carassius auratus*) and gudgeon (*Gobio gobio*); the Siluriformes, such as white catfish (*Ameiurus catus*) and wels catfish (*Silurus glanis*); and the Characiformes, such as red-bellied piranha (*Pygocentrus nattereri*) and Mexican tetra (*Astyanax mexicanus*; Dymek et al., 2020; Fishelson, 1997; Hara, 1975; Lazzari et al., 2014; Teichmann, 1954; Yamamoto & Ueda, 1979). More comparative data are clearly needed before we can understand the evolution of sensory systems in teleosts.

The findings of this study may have important methodological implications. Teleost fishes, in particular guppy and zebrafish, are becoming increasingly popular in cognition research and as model for investigation of cognitive deficits associated with human neuropathologies (Brown, Laland, & Krause, 2011; Stewart & Kalueff, 2012; Tierney, 2011). Cognitive studies in fish implicitly assume that all diurnal species are visually oriented and cognitive tasks most often require responding to visual stimuli, a practice that may penalise olfactorily oriented species. For example, zebrafish was found to perform poorly compared to other fish species in both numerical tasks and shape discriminations, an outcome that was attributed to cognitive differences (Agrillo, Miletto Petrazzini, Tagliapietra, & Bisazza, 2012; Gatto, Lucon-Xiccato, Bisazza, Manabe, & Dadda, 2020). In light of what we found in this study, it is instead possible that these differences are due to a different propensity to attend to stimuli that differ only in their visual features. Indeed, zebrafish have shown excellent learning, memory and discrimination capacities when tested with olfactory stimuli (e.g., Namekawa, Moenig, & Friedrich, 2018; Morin, de Souza Silva, Müller, Hardigan, & Spieler, 2013). Future research should examine in more depth how the sensory characteristics of the different species affect their performance in the extremely simplified laboratory situations.

## ACKNOWLEDGMENTS

We would like to thank Giacomo Federico Rubini, Marika Fanari and Andrea Ferrato for their help with testing the animals and with video analyses. This research was supported by PRIN 2015 Grant (Prot. 2015FFATB7) to A.B. from University of Padova. This work was carried out within the scope of the project “use-inspired basic research”, for which the Department of General Psychology of the University of Padova has been recognized as “Dipartimento di eccellenza” by the Ministry of University and Research (MIUR).

